# The phospholipid biosynthesis enzyme PlsB contains three distinct domains for membrane association, lysophosphatidic acid synthesis and dimerization

**DOI:** 10.1101/2023.04.30.538836

**Authors:** Yumei Li, Anjie Li, Zhenfeng Liu

## Abstract

Biosynthesis of phospholipids is fundamental for membrane biogenesis in all living organisms. As a member of the Glycerol-3-phosphate (G3P) Acyltransferase (GPAT) family, PlsB is a crucial enzyme catalyzing the first step of phospholipid synthesis by converting G3P and fatty acyl-coenzyme A (CoA)/acyl-carrier protein (ACP) into lysophosphatidic acid and free CoA (CoASH)/ACP. In bacterial cells, PlsB participates in the formation of persister cells related to multidrug tolerance, and is hence considered as a potential target for anti-persister therapy. By using the single-particle cryo-electron microscopy (cryo-EM) method, we have solved the structure of full-length PlsB from *Themomonas haemolytica* (*Th*PlsB) at 2.79 Å resolution. The *Th*PlsB protein forms a homodimer with *C*2 symmetry and each monomer contains three distinct domains, namely the amino-terminal domain (NTD), the middle catalytic domain (MCD) and the carboxy-terminal domain (CTD). For the first time, we have unraveled the binding sites of a fatty acyl-CoA and a 1,2-dioleoyl-sn-glycero-3-phosphate (DOPA) molecule in the MCD of PlsB. The interactions between *Th*PlsB and the membrane involve two surface-exposed amphipathic regions located in the NTD and MCD respectively. The results of structural and biochemical analyses suggest a membrane surface association-catalysis coupling model for the PlsB-mediated biosynthesis of lysophosphatidic acid occurring at the membrane-cytosol interface.

## Introduction

As the basic components of biological membranes, phospholipids are amphipathic molecules essential for all cellular organisms. Moreover, some phospholipids, such as phosphoinositide and phosphatidylserine, also function as messenger molecules for a diverse range of biological processes (Wang and Tontonoz, 2019). The initial steps of the *de novo* synthesis of phospholipids occur similarly in prokaryotes and eukaryotes. For instance, the Glycerol-3-phosphate Acyltransferase, known as PlsB in prokaryotes or GPAT in eukaryotes, catalyzes the acylation reaction at sn-1 position of G3P by using acyl-CoA or acyl-ACP as the donor of fatty acyl group (Lightner et al., 1980). There are two routes for lyso-phosphatidic acid (LPA) synthesis in bacteria, with the PlsB pathway being found mostly in proteobacteria and the PlsX/Y pathway carrying out a similar function in other bacteria (Kim et al., 2009; Lu et al., 2006). Recently, the structure of PlsY has been solved through X-ray crystallography, providing detailed insights into the molecular mechanism of LPA synthesis from acyl-phosphate and G3P (Li et al., 2017).

PlsB is a membrane-associated enzyme purified to high homogeneity from *E. coli* cells back in 1980 and exhibits the sn-glycerol-3-phosphate acyltransferase activity (Larson et al., 1980). It is the first bacterial acyltransferase characterized in detail (Zhang and Rock, 2008) and belongs to the superfamily of Acyl-CoA: glycerol-3-phosphate acyltransferase (GPAT) and acyl-CoA:1-acyl-glycerol-3-phosphate acyltransferase (AGPAT), which includes enzymes with four conserved motifs that are crucial for the acyltransferase function (Yamashita et al., 2014). The first conserved motif named Motif I has a characteristic HXXXXD (HX_4_D) sequence (Heath and Rock, 1998). It was demonstrated that the enzyme activity of PlsB from *E. coli* is almost completely lost when His306 or Asp311 in Motif I was mutated to Ala (Heath and Rock, 1998). Therefore, Motif I is essential for the catalytic activity of the GPAT enzymes (Dircks et al., 1999; Lewin et al., 1999).

Recently, PlsB has been shown to be the sole enzyme in bacteria that regulates steady-state cell membrane synthesis (Noga et al., 2020). It was also discovered that PlsB is involved in post-translational regulation of phospholipid synthesis and controls bacterial growth (Noga et al., 2020). The ability of bacteria to survive in different environments depends critically on the adjustment of their membrane lipid composition (Zhang and Rock, 2008). As PlsB is primarily responsible for controlling the synthesis of phospholipids during a steady state of cell growth, it is involved in maintaining the homeostasis of membrane lipids in bacteria. Moreover, PlsB may also participate in the regulation of lipopolysaccharide production indirectly during steady-state growth (Noga et al., 2020). Furthermore, PlsB plays a crucial role in the development of persister cells, which may become resistant to antibiotics. The expression of proteins involved in non-essential processes is downregulated in persister cells, turning them into inactive low-energy cells. Although persister cells have a non-heritable phenotypic tolerance to antibiotics, they can serve as an intermediate state to promote the formation of antibiotic-resistant bacteria (Spoering et al., 2006). Persister cells can result in infections that are resistant to multiple drugs, posing a significant negative impact on human health (Costerton et al., 1999; Lewis, 2001). As a glycerol-3-phosphate acyltransferase essential for bacterial persister cell formation, PlsB is considered as a potential target for the treatment of infection caused by persistent bacteria.

Eukaryotic PlsB homologs, namely GPATs, have been found in both animals and plants (Gimeno and Cao, 2008; Murata and Tasaka, 1997). In mammalian cells, there are many different isoforms of GPAT, and they function in different subcellular locations or within different tissues (Takeuchi and Reue, 2009). They also perform important physiological functions, such as regulating intracellular lipid accumulation, and have also been implicated to be related with diseases, such as obesity, insulin resistance and cancers (e.g., lung, breast, and prostate cancers, melanoma) (Ellis et al., 2012; Nagle et al., 2007; Nakagawa et al., 2012; Pellon-Maison et al., 2014). Although the crystal structure of a plant glycerol-3-phosphate acyltransferase has been solved (Turnbull et al., 2001), the overall structure of bacterial PlsB remains unknown. Little is known about the exact binding sites of acyl-CoA and G3P in PlsB and the mechanism of its association with membrane is enigmatic. Here, we describe the cryo-EM structures of PlsB from *Themomonas haemolytica* (*Th*PlsB), unravelling the substrate binding sites of PlsB and the potential connection between its enzymatic function and phospholipid interactions.

## Result

### Overall structure of *Themomonas haemolytica* PlsB

To understand the molecular basis underlying the biological process of LPA synthesis, we have solved the structure of PlsB from *Themomonas haemolytica* (*Th*PlsB) through the single-particle cryo-electron microscopy method. The final cryo-EM map, obtained through careful 2D classification, 3D classification, refinement and postprocessing procedure, reached an overall resolution of 2.79 Å (Supplementary Figure 1 and Table 1). *Th*PlsB forms an arch-shaped homodimer with a *C*2 symmetry. The *Th*PlsB dimer measures 198 Å in length and 80 Å in height, with a slightly curved shape, two surface grooves in the middle of the dimer and two lipid molecules associated with each monomer (Fig.1a-d). There is a hemispherical detergent micelle associated with each monomer on the concaved side, suggesting that *Th*PlsB protein may associate with membrane surface through the hydrophobic regions on the side buried by detergent micelles. As the detergent micelles only associate with *Th*PlsB on one side of its surface instead of forming a belt wrapping around the protein (Fig. 1a), the observation suggest that PlsB is most likely a monotopic membrane protein instead of a bitopic or polytopic membrane protein (Allen et al., 2019).

**Fig1. Overall.**
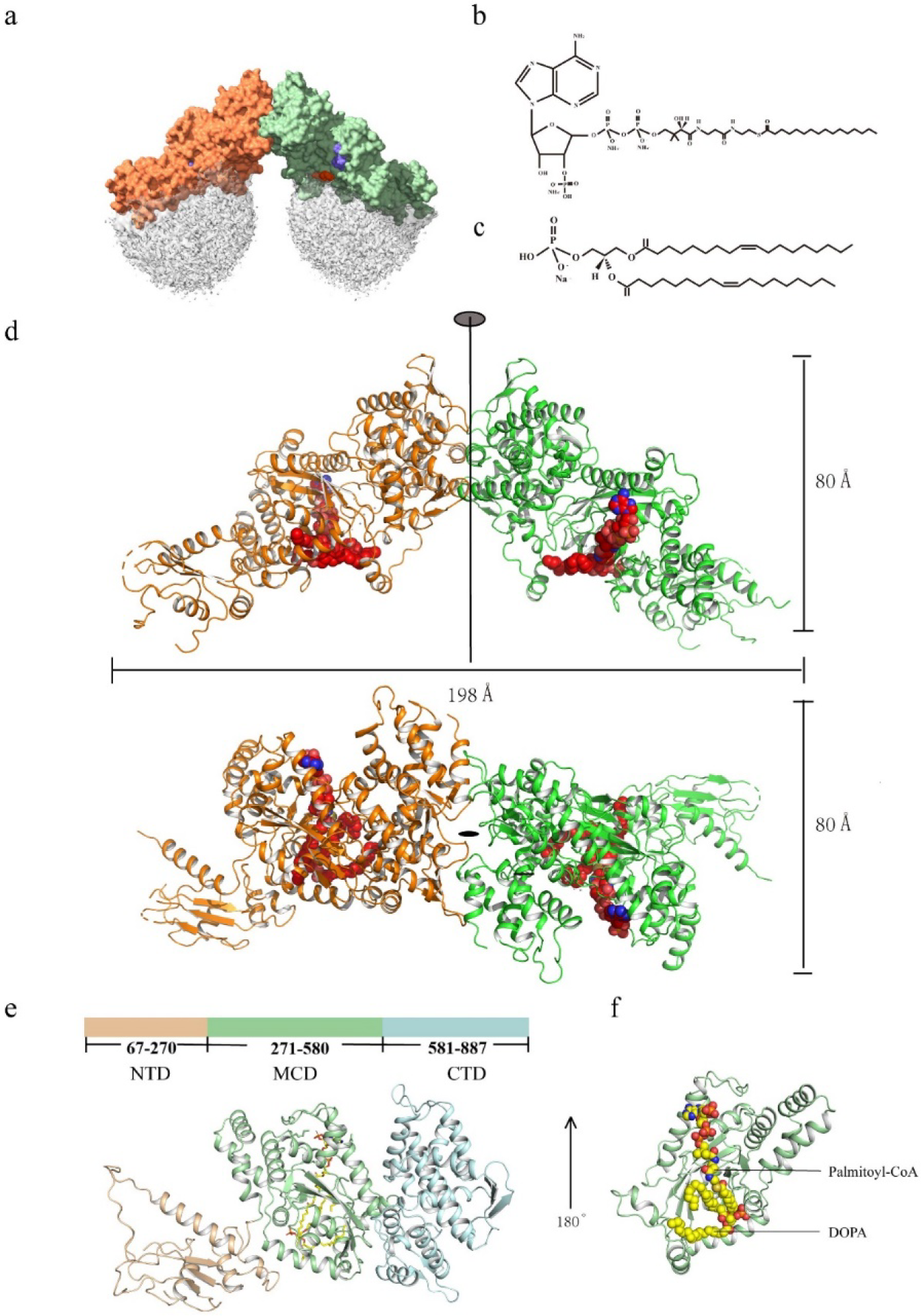
Structure of *Th*PlsB dimer and monomer. **a**, Cryo-EM density of *Th*PlsB dimer (in orange and green) anchored on the detergent micelles (in silver). **b**, Chemical structure of palmitoyl-CoA. **c**, Chemical structure of 1,2-dioleoyl-sn-glycero-3-phosphate (DOPA). **d**, The side view and top view of *Th*PlsB dimer. The proteins are presented as cartoon models, whereas the small molecules are shown as sphere models. **e**, Domain organization of a *Th*PlsB monomer. The NTD domain (1-270 aa), the MCD domain (271-580 aa) and the CTD domain are colored in golden, light green and cyan, respectively. **f**, The palmitoyl-CoA and DOPA molecules associated with the MCD of *Th*PlsB.

**Table 1.** Amino acid residues at the dimerization interface

| Monomer A | Monomer B | Distance (Å) | Bonds |
| --- | --- | --- | --- |
| Arg 866[NH1] | Asp 605[OD1] | 3.09 | Salt bridge/ Hydrogen bond |
| Arg 866[NH2] | Asp 605[OD2] | 3.75 | Salt bridge/ Hydrogen bond |
| Arg 866[NH1] | Asp 605[OD2] | 3.67 | Salt bridge |
| Arg 866[NH1] | Asp 759[OD1] | 2.71 | Salt bridge |
| His 870[NE2] | Asp 759[OD1] | 2.81 | Salt bridge |
| Arg866[ NH1] | Asp 759[OD1] | 3.76 | Salt bridge |
| His 870[NE2] | Asp 759[OD2] | 3.45 | Salt bridge |
| Asp 605[OD1] | Arg 866[NH1] | 2.88 | Salt bridge |
| Asp 605[OD2] | Arg 866[NH2] | 3.52 | Salt bridge |
| Asp 605[OD2] | Arg 866[NH1] | 3.22 | Salt bridge |
| Asp 759[OD1] | Arg 866[ NE ] | 3.99 | Salt bridge |
| Asp 759[OD1] | Arg 866[NH1] | 3.15 | Salt bridge |
| Asp 759[OD1] | His 870[NE2] | 2.93 | Salt bridge |
| Asp 759[OD2] | His 870[NE2] | 3.68 | Salt bridge |
| Lys 874[ NZ ] | Leu 653[ O ] | 3.76 | Hydrogen bond |
| Arg 808[ NE ] | Gly 654[ O ] | 2.85 | Hydrogen bond |
| Arg 808[ NH2] | Asp 655[ O ] | 3.73 | Hydrogen bond |
| Arg 869[ NH2] | Gln 757[ O ] | 3.01 | Hydrogen bond |
| Arg 866[ N ] | Gln 757[OE1] | 2.68 | Hydrogen bond |
| His 870[ NE2] | ASP 759[OD1] | 2.81 | Hydrogen bond |
| Lys 598[ NZ ] | Ser 810[ O ] | 3.44 | Hydrogen bond |
| Lys 598[ NZ ] | Tyr 813[ O ] | 2.53 | Hydrogen bond |
| Leu 653[ O ] | Lys 874[ NZ ] | 3.40 | Hydrogen bond |
| Gly 654[ O ] | Arg 808[ NE ] | 2.55 | Hydrogen bond |
| Asp 655[ O ] | Arg 808[NH2] | 3.38 | Hydrogen bond |
| Gln 757[ O ] | Arg 869[NH2] | 3.03 | Hydrogen bond |
| Gln 757[OE1] | Arg 866[ N ] | 2.77 | Hydrogen bond |
| Ser 810[ O ] | Lys 598[ NZ ] | 3.56 | Hydrogen bond |
| Tyr 813[ O ] | Lys 598[ NZ ] | 2.97 | Hydrogen bond |

Each *Th*PlsB monomer contains three distinct domains, namely the amino-terminal domain (NTD, 1-270), the middle catalytic domain (MCD, 271-580) and the carboxy-terminal domain (CTD, 581-877) (Fig. 1e). The NTD may have a crucial role in helping the *Th*PlsB protein to anchor on the membrane as its surface on the concaved side is largely buried in the detergent micelle. Meanwhile, the MCD may also be involved in membrane association as it contains a helix-loop motif buried in the detergent micelle. The MCD also harbors an acyltransferase αβ protein fold resembling that of PlsC protein (Robertson et al., 2017) and a HX_4_D (His366-Asp371) active site potentially involved in catalyzing the formation of LPA from glycerol-3-phosphate and acyl-CoA. Remarkably, there are two binding sites for small molecules in the MCD of each monomer (Fig. 1f), hosting a palmitoyl coenzyme A (palmitoyl-CoA) molecule (Fig.1b) and a 1,2-dioleoyl-sn-glycero-3-phosphate (dioleoyl phosphatidic acid, DOPA) molecule (Fig.1c) respectively. The palmitoyl-CoA and DOPA were added into the *Th*PlsB protein during sample preparation process. While one of the hydrophobic acyl chains of DOPA forms close interactions with palmitoyl-CoA, the polar phosphate head group binds to a peripheral site distant from the head group of palmitoyl-CoA.

The CTD forms a compact fold with fourteen α-helices in the core and two β-hairpins flanking on the sides. It contributes to dimerization of *Th*PlsB by interacting closely with the symmetry-related CTD from the adjacent monomer through three α-helices and six loops at the dimerization interface. There are 1560 Å^2^ surface area buried at the dimer interface and the amino acid residues involved in dimer formation are listed in Table 1.

### The substrate binding site of *Th*PlsB

Previously, it was demonstrated that PlsB can utilize either acyl-ACP or acyl-CoA to acylate the 1-position of glycerol-3-phosphate (Green et al., 1981). To locate the acyl-CoA binding site of PlsB, we have introduced palmityl-CoA (along with DOPA) to the purified *Th*PlsB protein during sample preparation. In a surface groove of the MCD in each *Th*PlsB monomer, there is an elongated small molecule density matching well with the model of palmitoyl-CoA (Fig. 2a). The characteristic bulky CoA head group is oriented towards the water-soluble side away from the detergent micelle, while the fatty acyl chain extends into a hydrophobic cleft facing the detergent micelle. The fatty acyl tail is wrapped around by hydrophobic amino acid residues, such as Tyr372, Leu373, Leu332, Val325 and Ile389. Meanwhile, the adenosine 3’,5’-diphosphate head group of palmitoyl-CoA form salt bridges or hydrogen bond with Arg414, Lys417, Lys458 and Lys558 (Fig.2b, c). Moreover, the 6-amine group of the adenine group form a hydrogen bond with Asp567.

**Figure 2.**
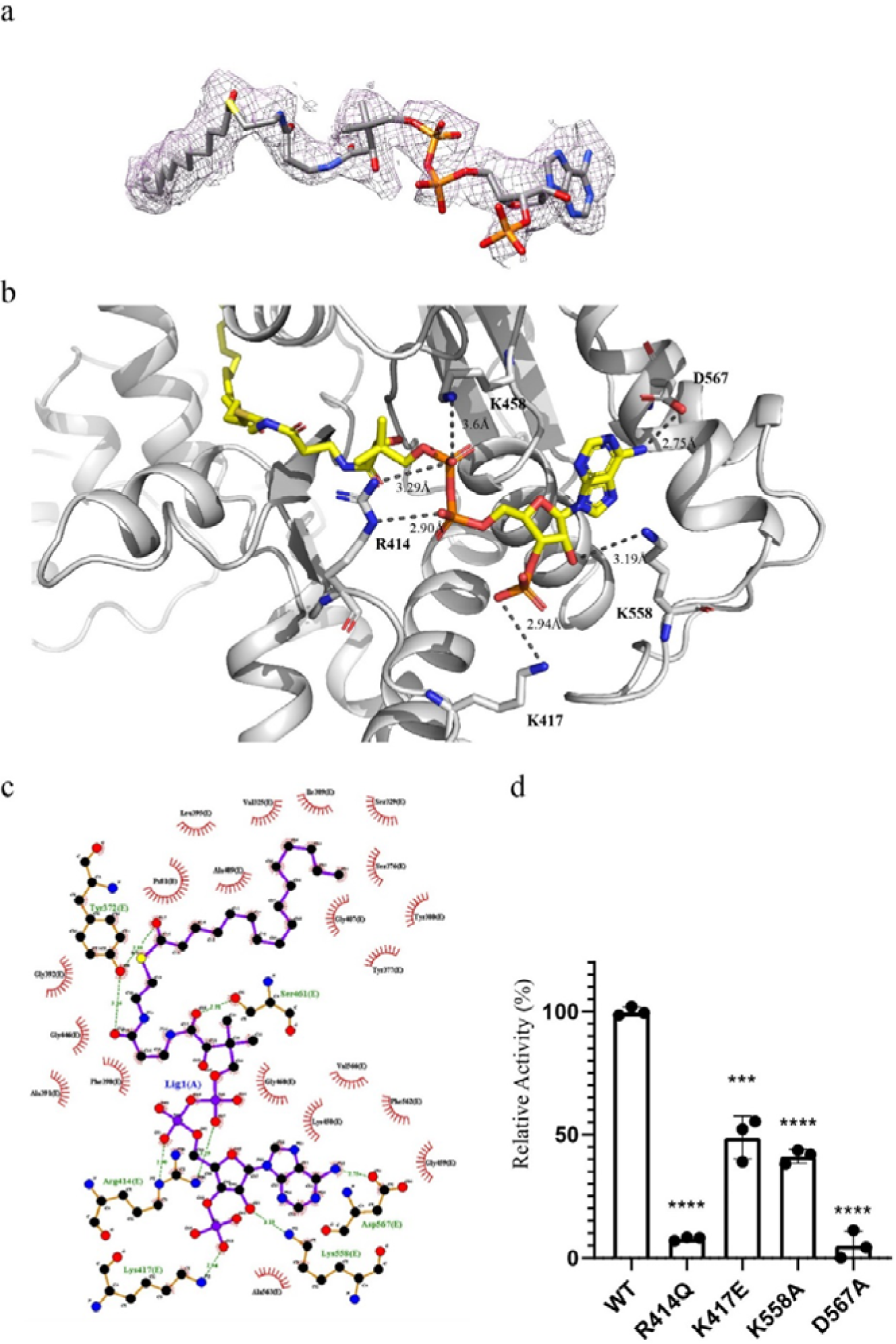
Substrate-binding site of *Th*PlsB. **a**, Cryo-EM density of the palmitoyl-CoA molecule associated with *Th*PlsB protein. The density is superposed with the stick model of palmitoyl-CoA. **b**, The palmitoyl-coenzyme A molecule forms salt bridges and a hydrogen bond with its nearby amino acids. **c**, The interactions between palmitoyl-CoA and nearby amino acids as revealed by Ligplot program. The green dashed lines indicate hydrogen bonds between the atoms involved, while the arc with spokes represent hydrophobic contacts between the ligand atoms and nearby groups. **d**, The enzymatic activities of four site-directed mutants of *Th*PlsB vs the wild type. ****, p≤0.0001; ***, 0.0001<p≤0.001 (p=0.0006). T test was used for the statistical analysis. The mean values ± standard deviation (SD, n=3) are presented.

To verify the functional role of the amino acid residues involved in binding the head group of palmitoyl-CoA, we have mutated some of the above amino acid residues and measured the enzyme activity of the mutants, such as R414Q, K417E, K558A and D567A. Compared with the wild type, the enzymatic activities of the mutants have been decreased by 48%-95%. Among them, the activities of R414Q and D567A mutants were only 10% and 5% of the wild type respectively, while the activities of K417E and K558A decreased to about 50% and 40% of the wild type (Fig. 2d). These substrate-binding sites are highly conserved among its homologs from various prokaryotic and eukaryotic species (Supplementary Fig. 2). While Arg414 and Tyr372 are highly conserved in homologs from both prokaryotes and eukaryotes, Lys417 is conserved in *E. coli* and *T. haemolytica*, but not in higher organisms. The data demonstrate that single point mutations in the amino acid residues interacting with palmitoyl-CoA will reduce the enzyme activities of *Th*PlsB protein dramatically. As revealed in the structure, they form specific interactions with the substrate molecule (Fig. 2b), and the mutations may affect the substrate-binding affinity and/or alter the position, orientation or conformation of the substrate molecule.

As shown in Fig. 3a, the His366 and Asp371 residues from the crucial HXXXXD motif at the active site do not form direct interactions with the palmitoyl-CoA molecule as their distances are too large. There might be unobserved water molecules filling in the gap between them, linking His366 and Asp371 with the substrate molecule. Mutations of the two central residues to Ala both lead to dramatic reduction of the activity of *Th*PlsB (Fig. 3b), consistent with the previous observation in *Ec*PlsB (Heath and Rock, 1998). Nearby His366, there is a second His residue (His369) located close to the palmitoyl-CoA molecule, and two Arg residues (Arg448 and Arg450) may potentially provide positively-charged side chains to attract and/or bind the negatively-charged phosphate group of the G3P molecule (Fig. 3c).

**Fig 3.**
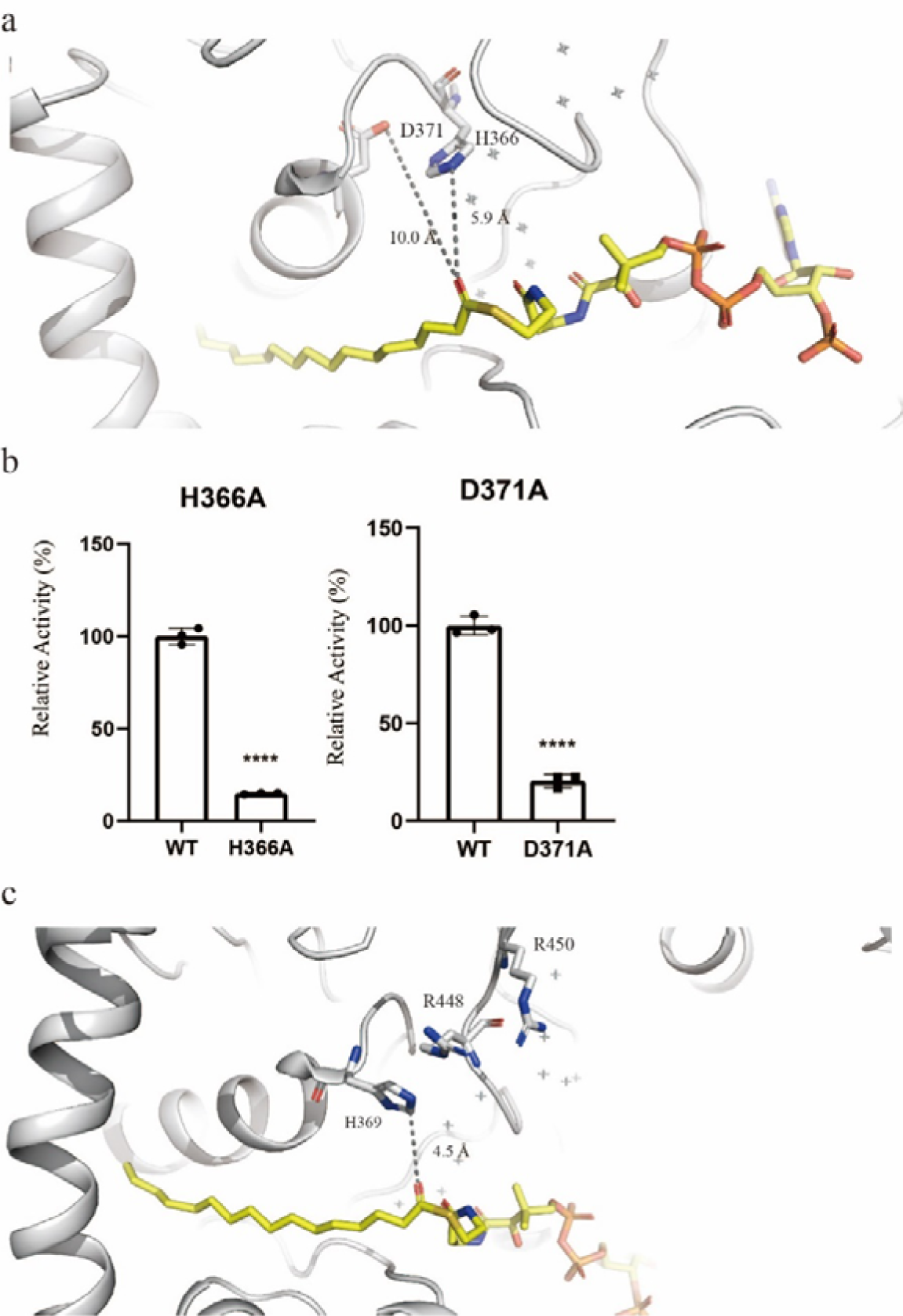
The active-site His366 and Asp371 residues in *Th*PlsB. **a**, The spatial relationships among His366, Asp371 and the palmitoyl-CoA. Note that His366 and Asp371 are the conserved amino acid residues in the HXXXXD motif characteristic of the GPAT/AGPAT family. **b**, The enzymatic activity of the H366A and D371A mutants of *Th*PlsB in comparison with the wild-type enzyme activity. ****, p≤0.0001. T test was used for the statistical analysis. The mean values ± standard deviation (SD, n=3) are presented. **c**, The distance and relative position between His369 (a second His residue nearby His366) and palmitoyl-CoA. Two positively-charged residues nearby His369, namely Arg448 and Arg450, are highlighted as stick models.

### Interaction of phospholipids with *Th*PlsB protein

The activity of PlsB from *E. coli* (*Ec*PlsB) is stimulated by various phospholipids, such as phosphatidylglycerol (PG), phosphatidylserine (PS), cardiolipin (CL) or phosphatidylethanolamine (PE) (Green et al., 1981). It was unclear whether there is a specific phospholipid-binding site in PlsB or how phospholipid binding affects the activity of PlsB. To address the question, we firstly investigated the interactions of *Th*PlsB with different phospholipids through the method of lipid blot assay. Different phospholipids were immobilized on the PVDF membranes, and the membranes were incubated with purified *Th*PlsB protein. After washing, the target protein associated with the membranes was probed by the antibody against the His-tag fused to *Th*PlsB. As shown in Fig. 4a, it is evident that *Th*PlsB associates strongly with CL and moderately with PA, but barely binds to PS, PG or PE.

**Fig 4.**
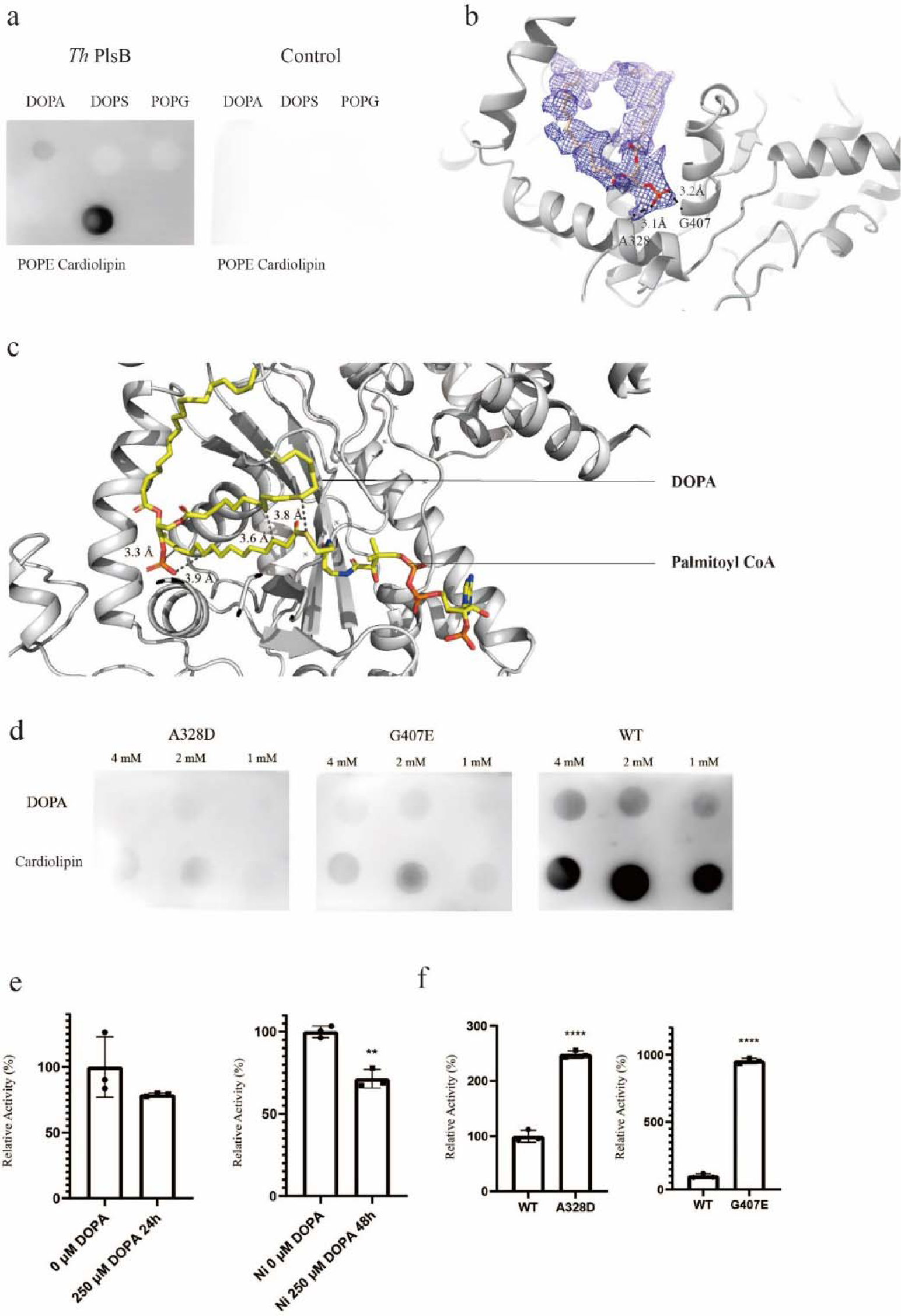
The binding site of DOPA in *Th*PlsB. **a**, Lipid blot analysis on the interactions between *Th*PlsB protein and various phospholipids. **b**, The location of a DOPA molecule in the structure of *Th*PlsB. The protein backbone is shown as silver cartoon model, while the cryo-EM density (blue meshes) of DOPA is superposed with the stick model. **c**. The relative positions of DOPA and palmitoyl coenzyme A molecules in the structure of *Th*PlsB. The dark dash lines indicate the potential van der Waals interactions between palmitoyl-CoA and DOPA. **d**, The interactions between A328D or G407E mutants and DOPA as well as cardiolipin (CL). The WT protein was loaded on a parallel blot for comparison with the mutants. **e**, The effect of DOPA on the enzymatic activity of *Th*PlsB. (left, the effect of 250 μM DOPA on *Th*PlsB activity after incubation for 24 h; right, the effect of 250 μM DOPA on *Th*PlsB activity after incubation for 48 h.) **f**, The enzymatic activity of G407E and A328D mutants in comparison with the WT.

The cryo-EM structure analysis revealed that there is a DOPA-like density in the map of *Th*PlsB (Fig. 4b). The putative DOPA molecule is located at the entrance of surface groove of MCD and the 1-acyl chain of DOPA interacts closely with the acyl group of palmitoyl-CoA molecule through hydrophobic interactions (Fig. 4c). The 2-acyl chain of DOPA extends along the hydrophobic surface of MCD lined by Met370, Met490, Val335, Ile339, Ile516 and Val519. Meanwhile, the phosphate head group of DOPA inserts in a small pocket surrounded by Gly407, Leu403, Leu404, V324 and Ala328, forming hydrogen bonds with the backbone amine of Gly407 and the carbonyl of Leu403 as well as van der Waals contacts with Val324 and Ala328.

To analyze the role of Gly407 and Ala328 in binding DOPA, we have mutated them to negatively charged amino acids. The G407E and A328D mutants exhibited much weaker binding to DOPA or CL than the wild type (Fig. 4d). In the presence of 250 μM DOPA, the activity of wild-type *Th*PlsB is slightly inhibited (Fig. 4e). DOPA is the product of the downstream enzyme PlsC utilizing LPA and acyl-CoA/acyl-ACP as substrates. Its inhibitory effect on PlsB activity may reflect a feedback regulation of phospholipid biosynthesis when DOPA is accumulated on the membrane at high concentration. For the G407E and A328D mutants with the DOPA-binding site affected, they exhibit much higher activities than the wild type (Fig. 4f). The negative charge introduced to the region around the active site may facilitate release of LPA from the active site and prevent the feedback inhibition by the product.

### Membrane-binding regions of *Th*PlsB

PlsB from *E. coli* is a membrane-associated enzyme as it could be extracted from membrane preparations with Triton X-100 (Larson et al., 1980). It was also suggested that PlsB may have two membrane-spanning domains (Zhang and Rock, 2008). The cryo-EM structure of *Th*PlsB indicates that the protein may have two membrane-anchoring domains (MAD1 and MAD2) instead of membrane-spanning domains, as the detergent micelles only cover part of the protein surface on one side rather than wrapping around them (Fig. 1a). The MAD1 and MAD2 of *Th*PlsB are located in the NTD and an α-helix-loop motif (Gly501-Ser522) region of the MCD, respectively (Fig. 5a).

**Fig 5.**
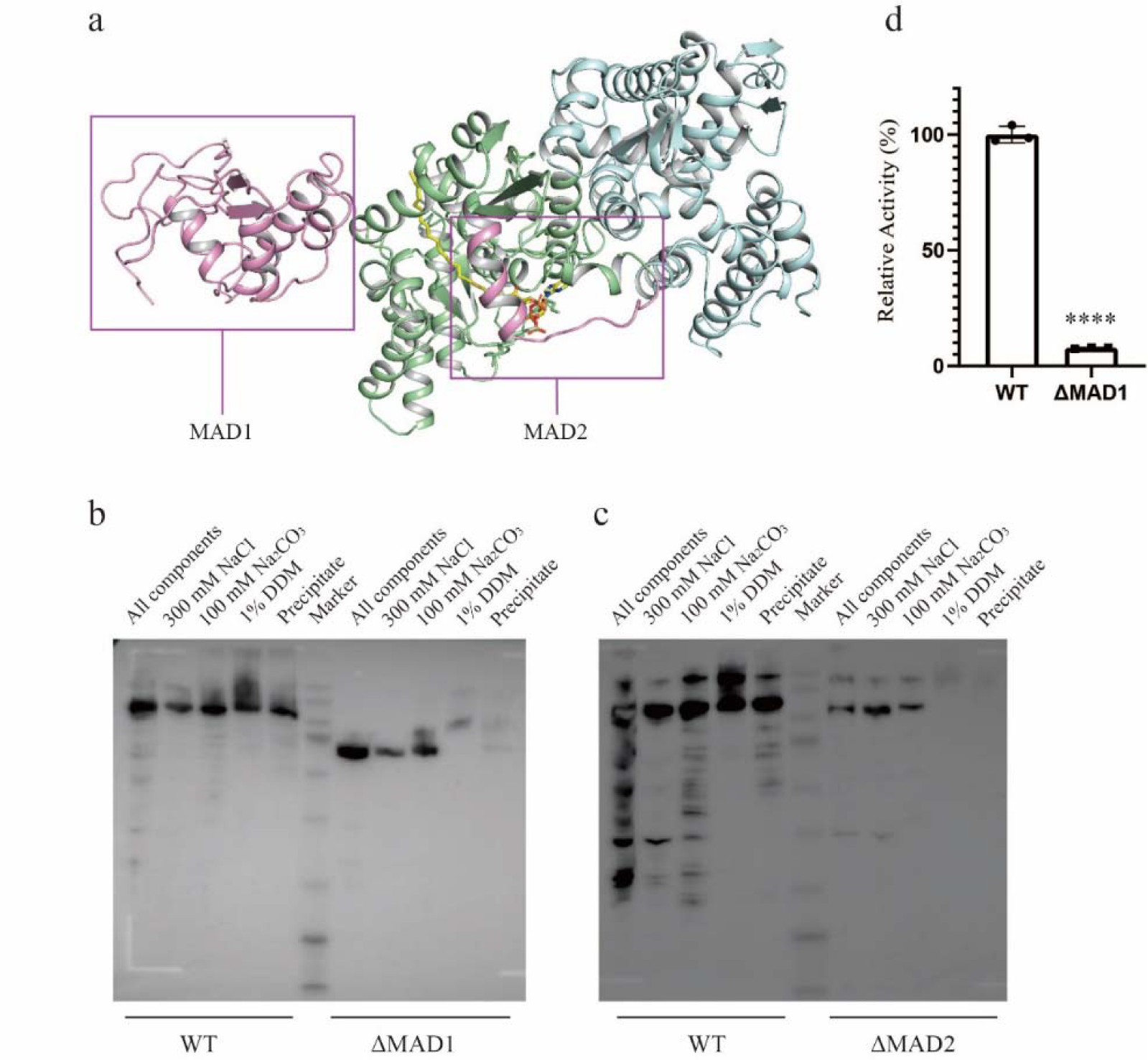
The membrane anchoring domains (MAD1 and MAD2) in *Th*PlsB. **a**, The MAD1 and MAD2 in *Th*PlsB monomer. The MAD1 and MAD2 in the boxed area are highlighted in pink, whereas the MCD and CTD are in green and cyan respectively. **b** and **c**, The membrane-washing/solubilization experiments for analyzing the interactions between ΔMAD1 (**b**)/ΔMAD2 (**c**) mutants (in comparison with the wild type/WT) and membrane. **d**, The enzymatic activity of ΔMAD1 mutant in comparison with that of WT.

While the MAD2 in MCD contains an amphipathic helix (Ser508-Arg521) with its hydrophobic residues buried in the micelle, the MAD1 in the NTD also contains a short amphipathic α-helix (Ala210-Asn220) covered by the micelle. As shown in Fig. 5b, the wild-type *Th*PlsB protein can be partially washed off the membrane when the membrane was treated with 100 mM Na_2_CO_3_ solution, or partially solubilized by 1% β-DDM. Therefore, *Th*PlsB is a peripheral membrane protein (or monotopic membrane protein) anchoring on the membrane instead of spanning the membrane. In contrast, the ΔMAD1 (Δ1-270) and ΔMAD2 (Δ501-520) mutants exhibit greatly reduced binding to the membrane, and most of the proteins can be washed off the membrane with 100 mM Na_2_CO_3_ (Fig. 5b and c). While the expression level of ΔMAD2 protein is much lower than the wild type, the ΔMAD1 protein expressed well in the *E. coli* cell. The activity of ΔMAD1 is only about 8% of the enzymatic activity of the wild type (Fig. 5d). Therefore, the close association of PlsB protein with the membrane is crucial for its function in catalyzing the synthesis of LPA. Truncation of MAD1 at the amino-terminal region not only affects the protein’s interaction with the membrane, but also reduces its enzymatic activity.

## Discussion

PlsB is a membrane-associated acyltransferase important for the *de novo* synthesis of phospholipids in γ-proteobacteria (Zhang and Rock, 2008). Eukaryotic homologs of PlsB, also known as GPAT, are present in various organisms including animals and plants (Zhang and Rock, 2008). Recent studies have demonstrated that PlsB is involved in the formation of persister cells in bacterial populations. PlsB is an essential and conserved bacterial persister protein that is a potential target for the development of drugs for treating infection caused by drug-resistant bacteria (Spoering et al., 2006). Furthermore, PlsB plays an important regulatory role in coordinating membrane synthesis with bacterial cell growth, while phospholipid synthesis is mainly regulated by PlsB and lipopolysaccharide synthesis may also be regulated through indirect control of PlsB (Noga et al., 2020). Recently, PlsB has been utilized for growing membranes under *in vitro* conditions in the field of synthetic biology research (Exterkate et al., 2018). Nevertheless, the structure of PlsB has remained elusive for decades and little is known about the mechanism of its function in catalyzing LPA synthesis on the membrane. The cryo-EM structure of *Th*PlsB serves as a framework for understanding the molecular basis of LPA production at the initial step of phospholipid biosynthesis, and may guide further optimization of the enzyme to enhance its activity for better application in the field of synthetic biology. As revealed in the cryo-EM structure, *Th*PlsB forms a symmetric homodimer in detergent solution. Previously, Gully et al. proposed that *Ec*PlsB may also exist in a dimeric state when they investigated protein interaction network for phospholipid synthesis on the *E. coli* membrane (Gully and Bouveret, 2006). In *Th*PlsB, the amino acid residues involved in dimerization of the protein are listed in Table 1. It is evident that dimerization of *Th*PlsB is mainly mediated by polar interactions, such as salt bridges and hydrogen bonds between adjacent monomers. The dimer forms an arch-shaped structure with a concaved surface on the side facing the membrane. When *Th*PlsB dimer associates with the membrane, the protein itself and the product LPA may induce deformation of the membrane in the local area in contact with the protein surface. LPA is an amphiphilic molecule having the potential to form positive spontaneous curvature of the membrane (Kooijman et al., 2005). The concaved surface of *Th*PlsB dimer on the side facing the membrane is likely adapted to a local convex membrane surface induced by the insertion of LPA to the membrane.

Through the lipid blot experiments, we have found that DOPA and cardiolipin interact with the *Th*PlsB protein. In the cryo-EM map of *Th*PlsB, we also found a lipid density nearby palmitoyl-CoA and assigned it as a DOPA molecule. One of the two fatty acyl tails of DOPA is nearly parallel to the acyl chain of the palmitoyl-CoA molecule and the phosphate head group is sandwiched between two amino acid residues, namely G407 and A328. The enzyme activity analysis results show that addition of 250 or 500 μM DOPA to the purified protein sample had inhibitory effect on its activity (Fig. 4e). Previously, it was demonstrated that phospholipids have a stimulating effect on the activity of *E. coli* PlsB (Green et al., 1981; Mishra and Kamisaka, 2001). The reason for the different effects of phospholipids on PlsB activity may be due to the species differences or the distinct experimental conditions between our work and the previous reports. Notably, the previous works used membrane fractions, whereas we used relatively more homogeneous protein samples purified in the detergent solution. Alternatively, it is also possible that DOPA and the other phospholipids have different binding sites on PlsB so that they could have distinct effects on the enzymatic activity. In either case, association of PlsB with the membrane surface is crucial for the access of phospholipid molecules from the membrane to the binding sites on PlsB in order for them to regulate the enzymatic activity.

The structure of *Th*PlsB presented in our work likely represents the acyl-CoA-bound and G3P-free state. Basing on the structural and biochemical insights obtained with *Th*PlsB, we propose a putative model to account for the mechanism of LPA synthesis catalyzed by PlsB (Figure 6). Assisted by the MAD1 in NTD and the MAD2 in MCD, PlsB is able to anchor on the cytoplasmic surface of plasma membrane. The substrates, namely acyl-coenzyme A and G3P may enter the active site cavity through the cleft between the MCD and CTD. The central conserved HX_4_D (His366-Asp371) motif is located on one side of palmitoyl-CoA-binding site. Asp371 is at 3.4 Å distance from Nδ of His366 forming a hydrogen bond and may help to maintain the lone pair of electrons in the Nε nitrogen in His366. Nearby His366, there is a blob of small molecule density (tentatively interpreted as two water molecules) surrounded by His369, Arg448, Arg450 and the β-mercaptoethylamine group of palmitoyl-CoA (Supplementary Fig. 3). The closest distance between the unidentified small molecule and the thiol sulfur atom of palmitoyl-CoA is only ∼4.5 Å. The site is favorable for binding of G3P as the phosphate group of G3P may be stabilized by the two Arg residues (Arg448 and Arg450) nearby and the 1-OH is positioned close to the thiol sulfur to launch nucleophilic attack on the thioester of palmitoyl-CoA. When the reaction is completed, the product LPA may enter the lipid bilayer and serves as the substrate for PlsC for the synthesis of PA, while the CoASH will be released to the cytoplasm (Figure. 6).

**Figure 6.**
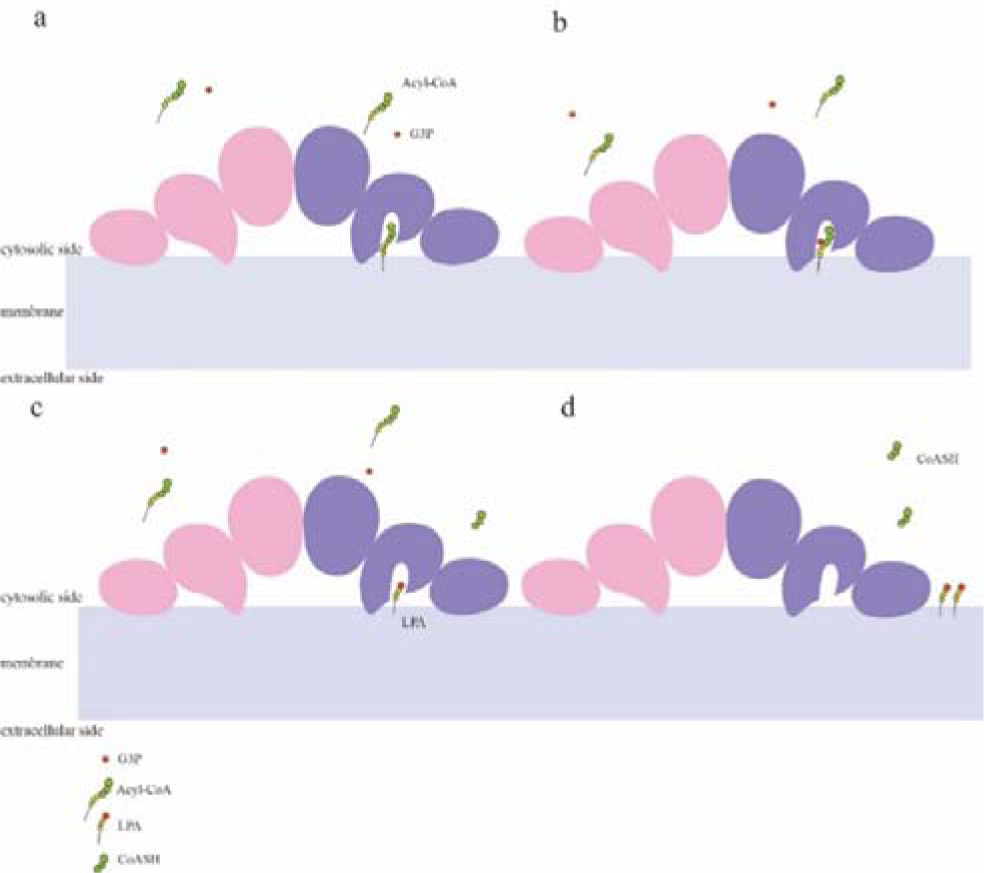
A putative cartoon model accounting for the catalytic mechanism of PlsB. **a**, Binding of acyl-CoA/acyl-ACP in the active-site cavity of MCD domain in PlsB. **b**, The intermediate state with both substrates loaded inside the active-site pocket. **c**, When the acyl-transfer reaction is completed, the free coenzyme A or ACP protein is released to the cytoplasm. **d**, The amphipathic product LPA likely diffuses into the membrane as a substrate for the downstream reaction.

## Method and Materials

### Protein expression and purification

The *Th*PlsB protein was expressed in *E coli* C41(DE3) cells by using pET 21b vector with synthesized cDNA (codon optimized through the GenSmart system from GenScript). The LB media with 0.1 mg mL^−1^ ampicillin were used for cell culture. To induce protein expression, 0.5 mM IPTG (Isopropyl-beta-D-thiogalactopyranoside) was added to the culture when the optical density OD_600_ reached 1.0. After further incubation at 37 °C for 2 h with constant shaking at 260 rpm, the cells were collected through centrifugation at 4000 ×g. The cell pellets were suspended in 10 volumes of Lysis Buffer (25 mM Tris-HCl, pH = 8.0, 300 mM KCl, 20 mM imidazole) and homogenized. To prevent protein degradation, 100 mM PMSF (phenylmethanesulfonyl fluoride) stock solution was added to the suspension to a final concentration of 1 mM. The cells were crushed by passing the homogenized suspension through a high-pressure homogenizer under 11,000 bar pressure for 3−5 rounds at 4 [. To solubilize membrane proteins, the detergent dodecyl-β-D-maltoside (β-DDM) was added to the cell lysate at a final concentration of 1% and incubated for 1.5 h at 4 [with constant stirring. The suspension was centrifuged at 18,000[rpm (JL-25.50 rotor, Beckman) for 30[m to remove insoluble matter. The supernatant was collected and loaded onto a Ni-NTA column, and it was washed by applying two different washing buffers (25 mM Tris-HCl, pH = 8.0, 300 mM KCl, 0.1 %GDN, 20 mM or 95 mM imidazole). Finally, a solution containing 300 mM imidazole was used for eluting the target protein, and the protein fractions were collected and concentrated to 4 mg/mL by using a 100 kDa molecular weight cut-off concentrator (Millipore).

The purified protein was incubated overnight with the substrate by adding palmitoyl coenzyme A at a molar ratio of 1:40 and phospholipid DOPA at 1:10 at 4[. The precipitate was removed from the solution by using high speed centrifugation (14000 ×g at 4°C for 10 min) and the supernatant was loaded onto a Superdex 200 increase 10/300 GL column. The buffer used for the size-exclusion chromatography is 25 mM Tris-HCl, pH = 8.0, 500 mM KCl, 2mM DTT, 5 mM EDTA and 0.1% GDN. The fractions around the peak at 9.7 mL were pooled and concentrated to 6 mg/mL using a 100 kDa cut-off concentrator.

### Cryo-EM sample preparation

The concentrated *Th*PlsB protein sample was applied onto the grids of Quantifoil-R1.2/1.3-Au-300 Mesh and blotted by using Vitrobot Mark IV which was maintained at 18°C with 80% humidity. In detail, the grids were glow discharged for 60s by using H_2_/O_2_, and 3µL of protein was loaded onto the grid, blotted for 3.5s with a force level of 2 and then waited for 5 s. The blotted grids were plunge frozen in liquid ethane and then transferred to liquid N_2_ for storage or data collection.

### Cryo-EM data collection and image processing

The frozen samples on EM grids were screened by using Talos F200C 200 kV microscope. The good ones were kept and used for data collection on Titan Krios 300 kV microscope with a Gatan K2 Summit direct electron detector camera and the SerialEM acquisition software. Energy filter was applied during data collection with a magnification of ×130,000. A total of 2,600 micrographs were collected at 1.04 Å/pixel, 60 e^−^ /Å^2^ with 40 movie frames and the defocus is at a range of −1.3 to −1.8μm.

Image processing was performed by using Relion 3.1 (Scheres, 2012; Zivanov et al. 2020). The micrographs were motion corrected by using the MotionCor2 program (Zheng et al., 2017) with 5 by 5 patches. Then CTFFIND4.1 (Rohou and Grigorieff, 2015) was used to for the estimation of the contrast transfer function (CTF) parameters. The misaligned or amorphous images were removed manually. Firstly, 1000 particles were picked manually and used for 2D Classification. Several good 2D class images were selected as references for further autopicking procedure. The autopicked particles were subject to multiple rounds of 2D Classification process. The 2D class images were selected and used for initial model building by using Cryosparc (Punjani et al., 2017). The model was then transferred to Relion to continue subsequent steps. All particles were divided into five 3D classes and further homogenous 3D refinement with the best 3D class of 37,575 particles yielded a 3D model at 2.91 Å resolution. Refinement with the other 3D classes did not give better resolution. To further improve the map quality, the second dataset was collected with 3,325 micrographs and was processed as in Supplementary Fig. 1b.

### Model building and refinement

For model building, predicted models of *Th*PlsB were generated by using the trRosetta server (Yang et al., 2020). The amino acid sequence of *Th*PlsB was input to the server and five possible models were subsequently obtained. The five predicted models were each superposed with the cryo-EM map in UCSF Chimera (Pettersen et al., 2021), and the one with the best match to the map was selected as the initial model for further adjustment and refinement.

The manual adjustment of the model was carried out in Coot (Emsley and Cowtan, 2004), starting from the MCD region as it fits well with the cryo-EM map. After the MCD model achieved optimal fitting with the densities, the CTD model was adjusted by referring the corresponding densities, followed by the NTD which exhibits relatively weaker densities. The position of each amino acid residue was manually adjusted so that it could have an optimal matching with the electron density map. After the model was adjusted manually, it was input to the Phenix program (Liebschner et al., 2019) for further refinement against the cryo-EM map. The first process of real-space refinement is simulated annealing, which helps to anneal the structure gradually from high to low temperature so as to find the minimal energy state. The refined structural model was adjusted manually by using Coot again. Finally, multiple rounds of minimization_global, local-grid-search and B-factor (adp) refinement are carried out to improve the structural model.

### Biochemical analysis on the association of *Th*PlsB protein with membrane

For membrane association analysis, 6 g of cells was suspended in 60 mL lysis buffer (25 mM Tris-HCl, pH = 8.0, 300 mM KCl, 20 mM imidazole) and the bacterial cells were lysed by using high pressure homogenizer as described above. After the lysate was centrifuged at 11000 ×g for 30 min, the supernatant was collected and used for ultracentrifugation at 36900 rpm for 45 m. The supernatant will be retained for SDS-PAGE sample preparation. The precipitate from the previous step was resuspended with the same volume of 100 mM Na_2_CO_3_ solution and the mixture was ultracentrifugated at 36,900 rpm for 45 m for the second time. The supernatant was also retained for SDS-PAGE sample preparation. The precipitate was suspended in the same volume of a solubilization solution (25 mM Tris-HCl 8.0, 300 mM NaCl, 1% β-DDM) and incubated at 4 for 1 h. After the sample was centrifuged at 36900 rpm for 45 m, the supernatant was collected and the insoluble pellets were also sampled for further analysis through SDS-PAGE. Subsequently, the samples taken from different stages of the experiment were mixed with 5 × SDS-PAGE loading buffer, and then loaded on the SDS-PAGE gel (12 %) for electrophoresis. Subsequently, western blot was performed to probe the target protein bands on the gel by using the Anti-His Mouse monoclonal antibody (1:80,000 dilution) and Goat anti-Mouse IgG (H + L)–HRP (1:20,000 dilution). The images of the western blots were taken by using the a chemiluminescence CCD system (ChemiScope 3500mini imager, Clinx Science Instruments).

### Thin layer chromatography and enzyme activity assay

We used a method combining TLC and isotope labeling to analyze the enzymatic activity of the *Th*PlsB protein. The purified wild-type and mutant *Th*PlsB proteins were diluted to 3 mg/ml for further activity-assay experiments. Initially, 5 uL protein sample was added into a buffer with 100 mM Tris-HCl 8.0, 1 mg/mL BSA and 50 mM palmitoyl CoA. 50 mM ^14^C-G3P is used as a labeled substrate which was added last and serves as the activator of the reaction. The total reaction volume was 50 mL. All reactions were proceeded at 37 for 2 h. As the catalytic reaction of the *Th*PlsB protein was fairly slow, we used 2 h as the reaction time to accumulate sufficient labeled products. The reactions were stopped by adding trichloroacetic acid (TCA) to the mixture at a final concentration of 0.5% (w/v). The mixture was centrifuged 10 m and 2 μL of the supernatant was applied onto the TLC aluminum plates. After the droplets were completely dry, the TLC plates were placed in a developing solvent (chloroform: methanol: acetic acid: water= 7.0: 1.8: 0.9: 0.30). Afterwards, the plates were pressed against the phosphorous screen for 3 days. The fluorescence image on the phosphorous screen was taken by using FLA-7000 (GE). The quantitative area analysis on the image was performed by using the Image J and GraphPad Prism 8 softwares.

### Lipid blot assay

The standard samples of phospholipids, including DOPA, DOPS, POPG, POPE, and cardiolipin, were obtained by dissolving the dry lipid powder in chloroform at a concentration of 4 mM. The PVDF films were cut into rectangular pieces of suitable size and loaded with spots of various phospholipids. After the lipid spots were dried, the film was then placed in a solution (1 ×PBS) containing 0.3% BSA. After being incubated for 2 hours at 25 °C, the films become nearly transparent. Subsequently, 10 μL of *Th*PlsB (1 mg/mL) was added to the mixture and incubated with the film overnight at 4 °C. The film was then washed with Phosphate buffered saline with Tween 20 (PBST) three times (15 minute each time). The protein associated with the lipid spots was probed by using the Anti-His Mouse monoclonal antibody and Goat anti-Mouse IgG (H + L)–HRP. The chemiluminescence CCD system (ChemiScope 3500mini imager, Clinx Science Instruments) was used for visualizing and imaging the blots.

## Author contributions

Yumei Li, Investigation (biochemical experiments, data collection and processing, model building and refinement), Data curation, Visualization, Writing the original draft and revision; Anjie Li; Investigation (data collection and processing); Zhenfeng Liu, Conceptualization, Methodology, Resources, Model building, Data curation, Writing, review & editing of the manuscript, Visualization, Supervision, Project administration, Funding acquisition.

## Competing interests

Authors declare no competing interests.

## Acknowledgements

We thank Bo-Ling Zhu, Xiao-Jun Huang, De-Yin Fan and other staff members for their support in cryo-EM data collection at the Center for Biological Imaging (CBI) in the Core Facilities for Protein Science (CFPS) at the Institute of Biophysics (IBP), Chinese Academy of Sciences, and Hong-jie Zhang at the Structural and Functional Analysis Laboratory in the CFPS, IBP for his assistance with enzyme activity assay. We are grateful to Dr. Dan-feng Song for his help with molecular dynamics simulation, Jian-Yu Shan for his support on figure preparation and Ms. Xiao-Bo Liang for her assistance in sample preparation and data storage. The project is funded by the Chinese Academy of Sciences (the Strategic Priority Research Program XDB37020101 and Young Scientists in Basic Research YSBR-015 to Z. L.) and the National Natural Science Foundation of China (31925024 to Z.L.).

## Notes

### Competing Interest Statement

The authors have declared no competing interest.

