## Supplementary Figures 1-3 and Table 1 for "The phospholipid biosynthesis enzyme PlsB contains three distinct domains for membrane association, lysophosphatidic acid synthesis and dimerization"

**Supplementary Data (Supplementary Figures 1-3 and Table 1)**

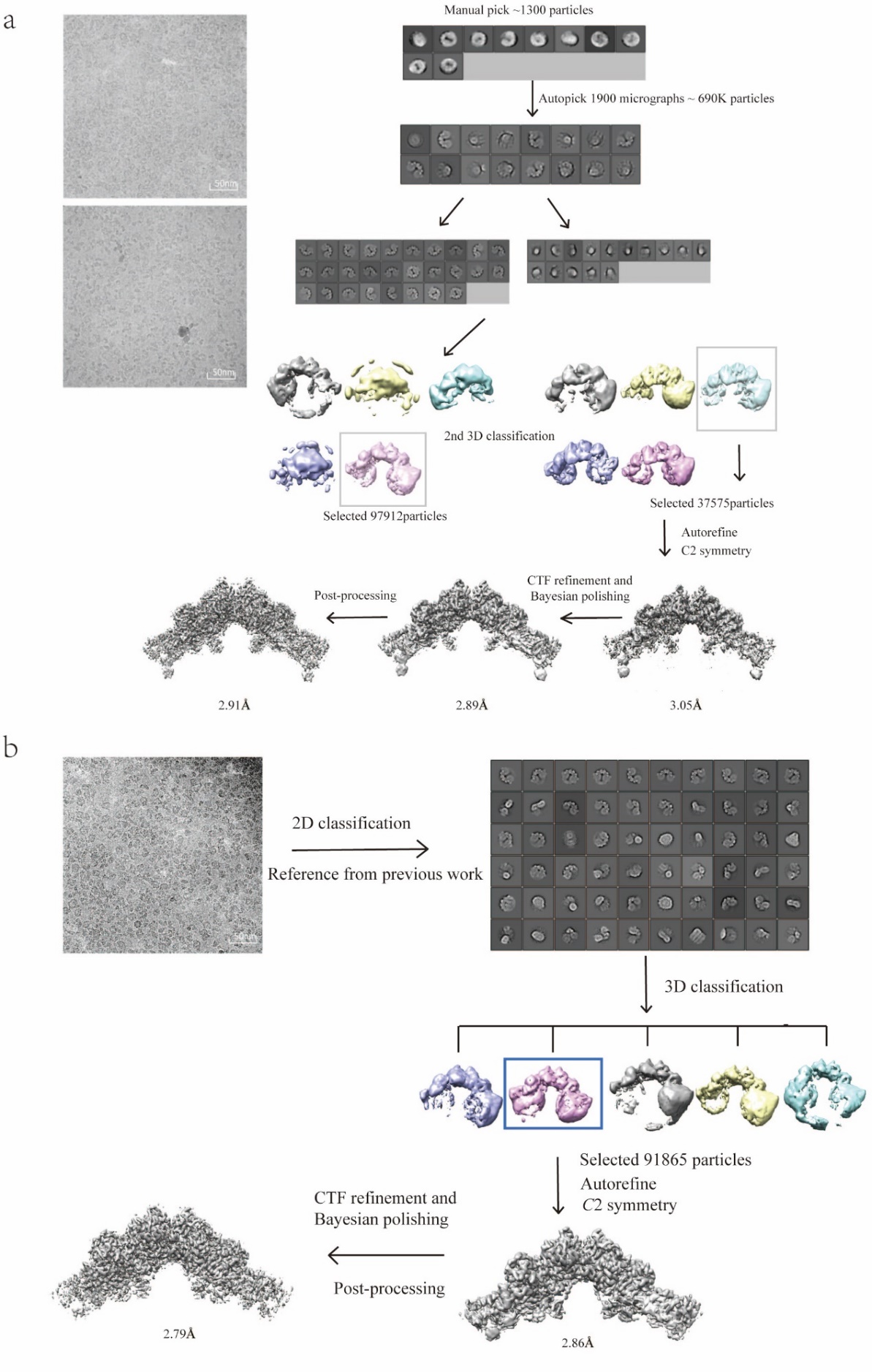

**Supplementary Figure 1.** Cryo-EM data processing and refinement. **a**, The overall scheme of cryo-EM data collection and the first-round processing and refinement. **b**, The scheme for the second-round data processing and refinement. The particles used to reconstruct the final map in work **a** were subjected to a further round of 2D classification, and the resulting 2D class averages were used as templates to guide the auto-picking in this dataset.

­

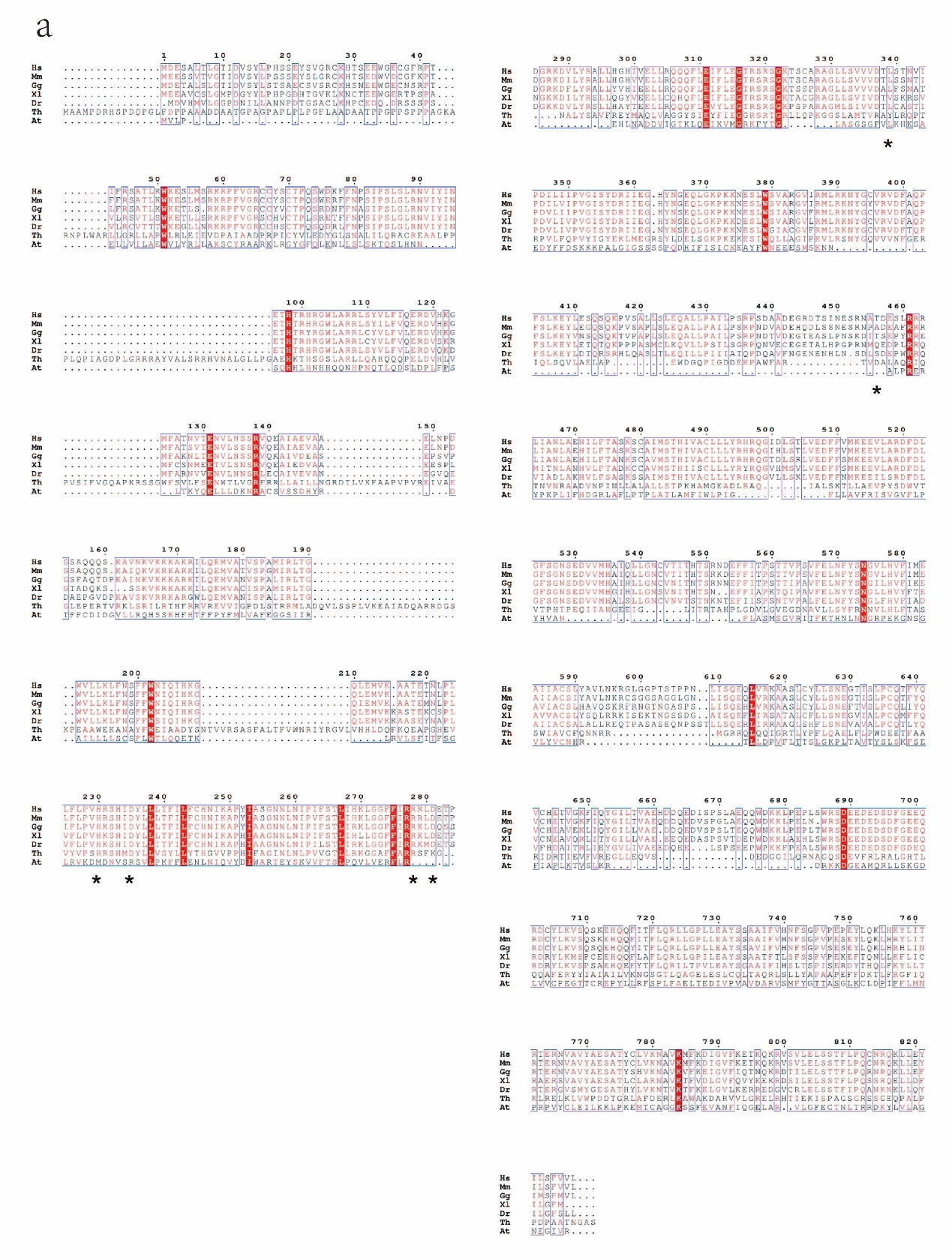

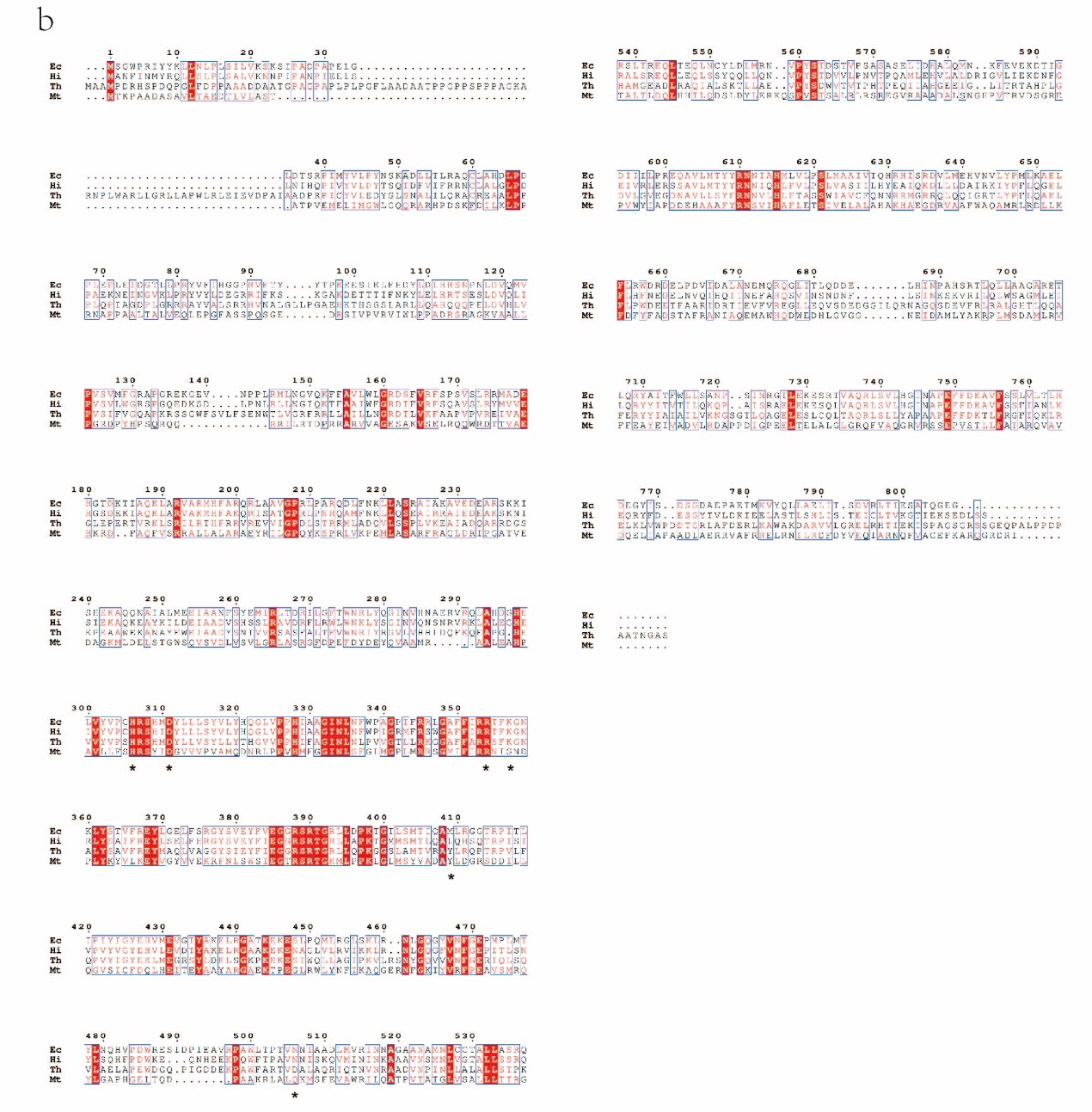

**Supplementary Figure 2. a**, Sequence alignment of *Th*PlsB and various eukaryotic GPAT homologs. Hs, *Homo sapiens;* Ms*, Mus musculus*; Gg*, Gallus gallus*; Xl*, Xenopus laevis*; Dr*, Danio rerio*; Th, *Themomonas haemolytica*. **b**, Sequence alignment diagram of *Th*PlsB and other prokaryotic PlsB orthologs. Ec, *Escherichia Coli;* Hi, *Haemophilus influenzae*; Mt, *Mycobacterium tuberculosis*. ­

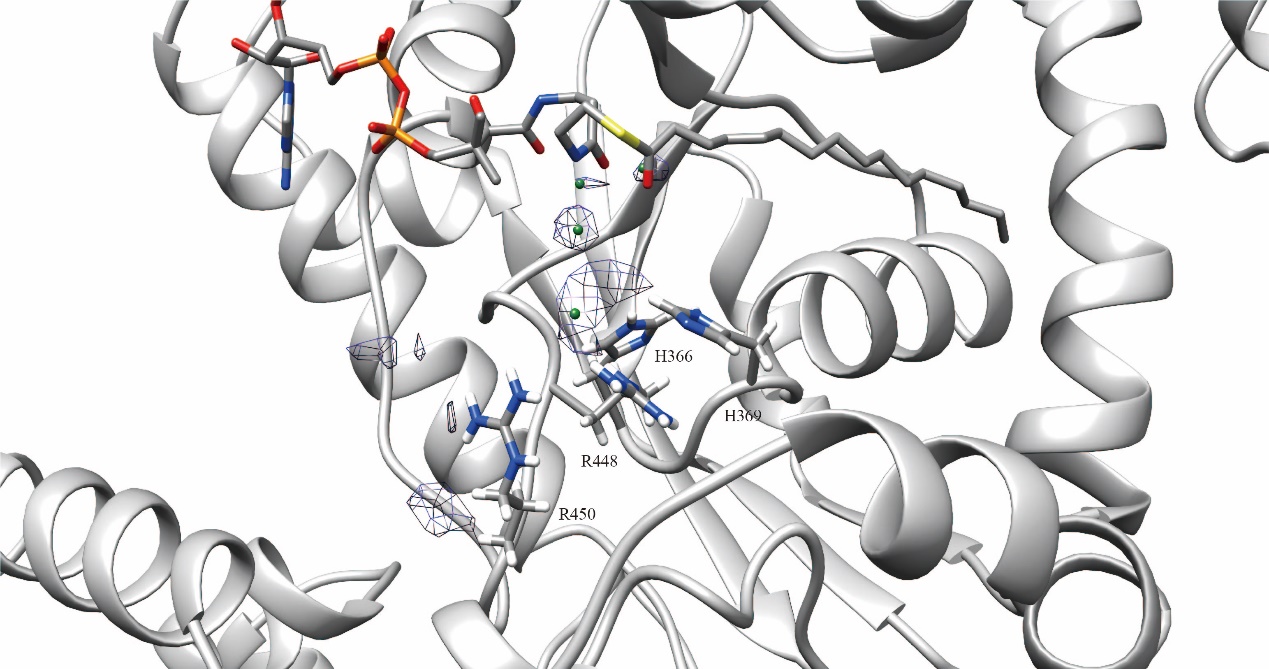

**Supplementary Figure 3. Two putative water molecules in the active site and their nearby amino acid residues.** The blue meshes in the middle region are the cryo-EM densities for potential water molecules (modeled as green spheres) in the active site. The water molecule-binding site may also serve as the binding site for a G3P molecule.

**Supplementary Table 1 Statistics of cryo-EM data collection, refinement and validation for *Th*PlsB structure.**

|  | *Th*PlsB |
| --- | --- |
| **Data collection and processing** |  |
| Magnifcation | 130,000 |
| Voltage (kV) | 300 |
| Electron exposure (e^-^/Å) | 60 |
| Defocus range (μm) | -1.3 ~ -1.8 |
| Pixel size (Å) | 1.04 |
| Symmetry imposed | *C*2 |
| Final particle images (no.) | 91,865 |
| FSC threshold | 0.143 |
| Map resolution (Å) | 2.79 |
| **Refinement**  Model resolution (Å)  FSC threshold  Map sharpening B factor (Å^2^) | 0.5  -95.0 |
| Model composition |  |
| Nonhydrogen atoms | 23,720 |
| Protein residues | 1,468 |
| Ligands | 4 |
| B factor (Å^2^)  Protein | 45.92 |
| Ligand | 37.94 |
| R.m.s. deviations |  |
| Bond lengths (Å) | 0.006 |
| Bond angles (°) | 0.905 |
| Validation |  |
| MolProbity score | 1.44 |
| Clash score | 2.91 |
| Poor rotamers (%) | 0.00 |
| Ramachandran plot |  |
| Favored (%) | 94.83 |
| Allowed (%) | 5.17 |
| Outliers (%) | 0.00 |
